# Functional Analysis of All Salmonid Genomes (FAASG): an international initiative supporting future salmonid research, conservation and aquaculture

**DOI:** 10.1101/095737

**Authors:** Daniel J. Macqueen, Craig R. Primmer, Ross D. Houston, Barbara F. Nowak, Louis Bernatchez, Steinar Bergseth, William S. Davidson, Cristian Gallardo-Escárate, Tom Goldammer, Yann Guiguen, Patricia Iturra, James W. Kijas, Ben F. Koop, Sigbjørn Lien, Alejandro Maass, Samuel A.M. Martin, Philip McGinnity, Martin Montecino, Kerry A. Naish, Krista M. Nichols, Kristinn Ólafsson, Stig W. Omholt, Yniv Palti, Graham S. Plastow, Caird E. Rexroad, Matthew L. Rise, Rachael J. Ritchie, Simen R. Sandve, Patricia M. Schulte, Alfredo Tello, Rodrigo Vidal, Jon Olav Vik, Anna Wargelius, José Manuel Yáñez

**Affiliations:** Institute of Biological and Environmental Sciences, University of Aberdeen, Aberdeen AB24 2TZ, United Kingdom.; Department of Biology, University of Turku, 20014, Finland.; The Roslin Institute and Royal (Dick) School of Veterinary Studies, The University of Edinburgh, Midlothian, EH25 9RG, UK.; Institute for Marine and Antarctic Studies, University of Tasmania Launceston, Tasmania, Australia.; Département de biologie, Institut de Biologie Intégrative et des Systèmes (IBIS), Université Laval, Québec, Canada, G1V 0A6.; The Research Council of Norway, Drammensveien 288, P.O.Box 564, NO-1327 Lysaker, Norway.; Department of Molecular Biology and Biochemistry, Simon Fraser University, Burnaby, British Columbia V5A 1S6, Canada.; Laboratory of Biotechnology and Aquatic Genomics, Interdisciplinary Center for Aquaculture Research, Department of Oceanography, Universidad de Concepción, Concepción 4030000, Chile; Leibniz Institute for Farm Animal Biology, Institute for Genome Biology, Fish Genetics Unit, Wilhelm-Stahl-Allee 2, 18196, Dummerstorf, Germany.; INRA, UR1037 Fish Physiology and Genomics, Rennes, France.; Faculty of Medicine, University of Chile, Santiago 8380453, Chile.; CSIRO Agriculture, St Lucia, QLD, 4067, Australia.; Department of Biology, University of Victoria, Victoria, British Columbia V8W 3N5, Canada.; Centre for Integrative Genetics (CIGENE), Department of Animal and Aquacultural Sciences, Norwegian University of Life Sciences, Ås NO-1432, Norway.; Center for Mathematical Modelling, Department of Mathematical Engineering, University of Chile, Santiago 8370456, Chile.; Center for Genome Regulation, University of Chile, Santiago 8370456, Chile.; School of Biological, Earth and Environmental Sciences, University College Cork, Cork, Ireland.; Center for Biomedical Research, Universidad Andres Bello, Santiago 8370146, Chile; FONDAP Center for Genome Regulation, Faculty of Biological Sciences and Faculty of Medicine, Universidad Andres Bello, Santiago 8370146.; School of Aquatic and Fishery Sciences, University of Washington, Box 355020, Seattle, WA 98195, USA.; Conservation Biology Division, Northwest Fisheries Science Center, National Marine Fisheries Service, National Oceanic and Atmospheric Administration, 2725 Montlake Blvd E, Seattle, WA, 98112, USA.; Matis Ltd, Vínlandsleið 12, 113 Reykjavík, Iceland.; NTNU - Norwegian University of Science and Technology, NO-7491 Trondheim, Norway.; National Center for Cool and Cold Water Aquaculture, UsDA ARS, 11861 Leetown Road, Kearneysville, WV, 25430, USA.; Department of Agricultural, Food, and Nutritional Science, University of Alberta, Edmonton, AB, Canada.; Office of National Programs, USDA ARS, 5601 Sunnyside Avenue, Beltsville, Maryland 20705-5148.; Department of Ocean Sciences, Memorial University of Newfoundland, 1 Marine Lab Road, St. John’s, NL, A1C 5s7, Canada.; Genome British Columbia, Suite 400 – 575, West 8th Avenue, Vancouver BC V5Z 0C4, Canada.; Department of Zoology, University of British Columbia, 6270 University Blvd., Vancouver, BC, V6T 1Z4, Canada.; Instituto Tecnológico del Salmón S.A., INTESAL de SalmonChile, Chile.; Laboratory of Molecular Ecology, Genomics, and Evolutionary Studies, Department of Biology, University of Santiago, Santiago 9170022, Chile.; Institute of Marine Research, P.O. Box 1870, Nordnes, NO-5817 Bergen, Norway.; Faculty of Veterinary and Animal Sciences, University of Chile, Av. Santa Rosa 11735, Santiago, Chile & Aquainnovo, Cardonal s/n, Puerto Montt, Chile.

**Keywords:** Salmonid fish, Genome function, Comparative approach, Phenotype & data standardization, Evolution & Ecology, Physiology, Aquaculture, Whole genome duplication, Data sharing, Open access

## Abstract

We describe an emerging initiative - the ‘Functional Analysis of All Salmonid Genomes’ (FAASG), which will leverage the extensive trait diversity that has evolved since a whole genome duplication event in the salmonid ancestor, to develop an integrative understanding of the functional genomic basis of phenotypic variation. The outcomes of FAASG will have diverse applications, ranging from improved understanding of genome evolution, through to improving the efficiency and sustainability of aquaculture production, supporting the future of fundamental and applied research in an iconic fish lineage of major societal importance.

## 1. The importance of salmonid fish: from evolution to sustainable food production

Salmonids have combined scientific, societal and economic importance that is unique among fish (reviewed in [1]). They are naturally distributed in fresh and marine habitats throughout the Northern hemisphere and have been introduced to South America, Australia, Africa and the Middle East. They perform key ecological functions, e.g. [2], but many populations are declining, and extensive effort is being directed towards their conservation and management, especially with respect to anthropogenic-driven change, e.g. [3]. Salmonids include at least 70 species (but are sometimes classified as > 200), possessing a rich diversity of adaptations and life-history strategies [4]. The great phenotypic diversity amongst salmonids provides an excellent study system to understand adaptive divergence and ecological speciation [4, 5] and was potentially facilitated by a whole genome duplication (WGD) in their common ancestor ~95 Mya [6, 7]. Salmonid aquaculture and capture fisheries (mainly of Atlantic salmon and *Oncorhynchus* species) play an important role in the economic and/or food security of several nations, accounting for 7.2 / 16.6% of all traded fish in terms of share by weight / value [8].

## 2. Rationale for the FAASG initiative

The FAASG initiative follows the recent publication of the genomes of Atlantic salmon (*S. salar*) [9] and rainbow trout (*O. mykiss*) [10], which have proved invaluable to salmonid researchers (section 3) and establish a solid foundation for generating reference genome sequences for other salmonid species (Fig. 1). The next step for salmonid research is to annotate genome function, considering species and populations of major scientific interest (sections 6, 7). This will lay foundations to understand how genotypes are translated to phenotypes via different layers of regulation of gene and protein expression. Covering a broad diversity of research in salmonid biology will aid this action and is best achieved by involving the broadest possible research community (section 9). FAASG will follow principles established by the ‘Functional Annotation of Animal Genomes’ (FAANG) consortium (section 5) [11], using standardized approaches for functional annotation, including bioinformatics protocols and pipelines exploiting knowledge from other species and through an array of experimental assays (Table 1, section 7). However, the FAASG framework (section 6, Fig. 1) will also exploit unique features of salmonid biology, including recent WGD and extensive phenotypic variation at both macro- and micro-evolutionary timescales, to generate broad mechanistic insights into genome evolution and adaptation.

**Figure 1.**
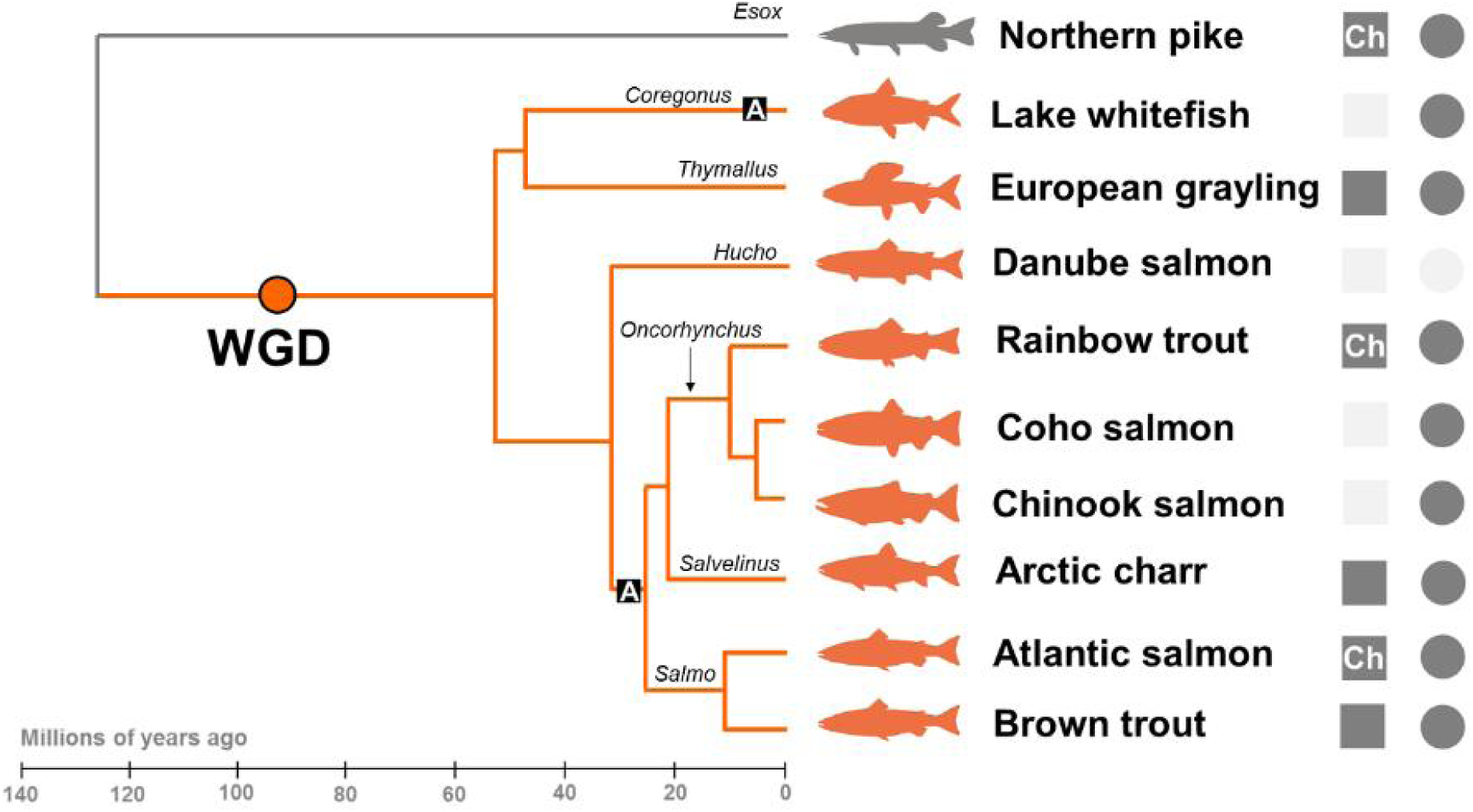
The comparative-evolutionary framework of FAASG. Shown are the main target lineages for functional annotation (see Table 1) and their evolutionary relationships (time-calibrated tree after [7]). The selected species come from all three salmonid subfamilies. The position of the salmonid-specific WGD is highlighted (after [7, 9–10]), along with Latin names of genera. Additional salmonid species that are future potential targets for functional annotation are not shown. Two lineages where anadromous life-history evolved are highlighted ‘A’ (after [47]). Relevant ‘omics resources are shown to the right of the tree - finalized (dark grey) or under active development (light grey), with ‘Ch’ highlighting chromosome-anchored genome assemblies.

**Table 1.**
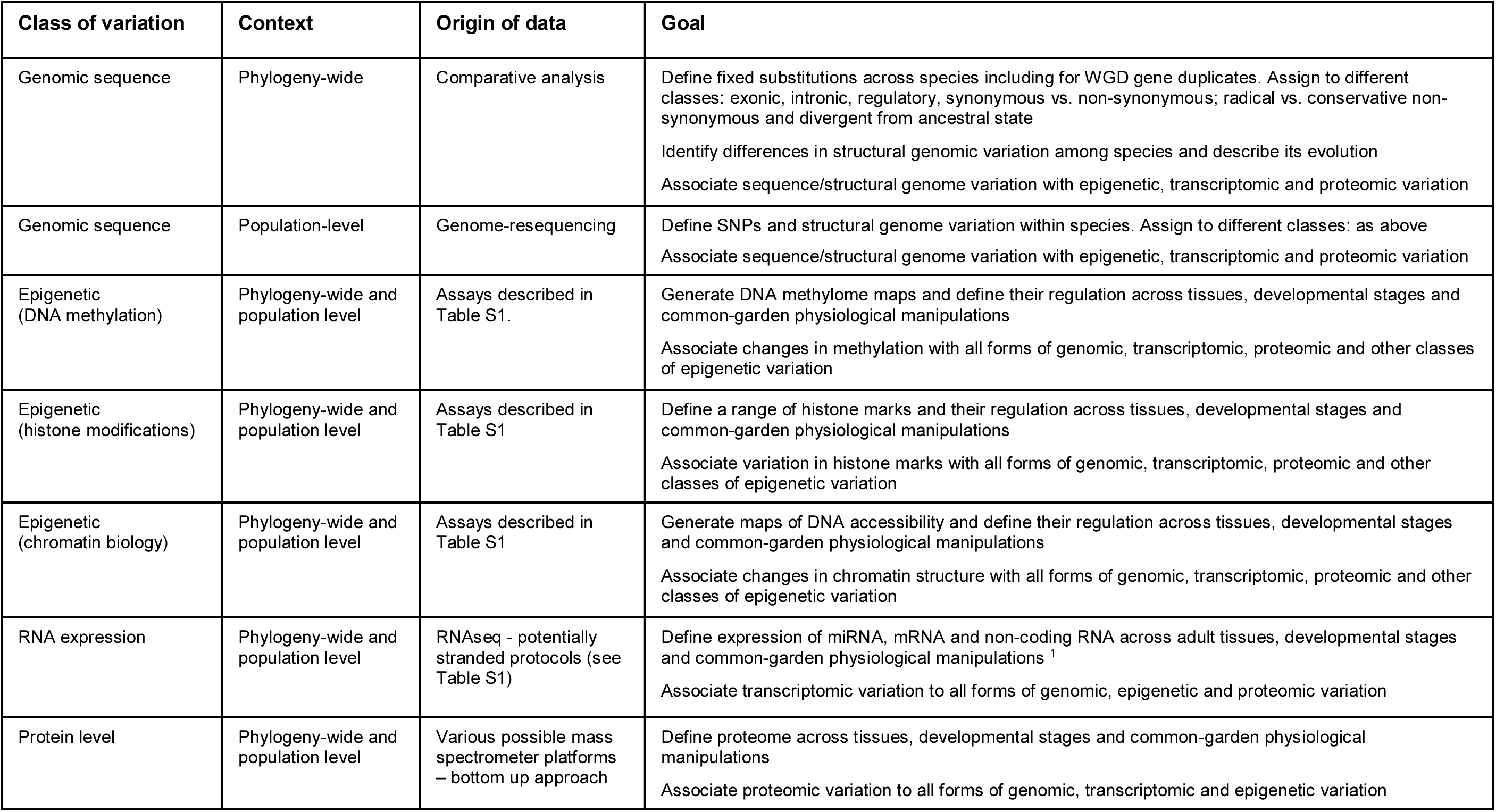
Levels of genome-wide functional annotation within the FAASG framework

| Class of variation | Context | Origin of data | Goal |
| --- | --- | --- | --- |
| Genomic sequence | Phylogeny-wide | Comparative analysis | Define fixed substitutions across species including for WGD gene duplicates. Assign to different classes: exonic, intronic, regulatory, synonymous vs. non-synonymous; radical vs. conservative non-synonymous and divergent from ancestral state<br>Identify differences in structural genomic variation among species and describe its evolution<br>Associate sequence/structural genome variation with epigenetic, transcriptomic and proteomic variation |
| Genomic sequence | Population-level | Genome-resequencing | Define SNPs and structural genome variation within species. Assign to different classes: as above<br>Associate sequence/structural genome variation with epigenetic, transcriptomic and proteomic variation |
| Epigenetic (DNA methylation) | Phylogeny-wide and population level | Assays described in Table S1. | Generate DNA methylome maps and define their regulation across tissues, developmental stages and common-garden physiological manipulations<br>Associate changes in methylation with all forms of genomic, transcriptomic, proteomic and other classes of epigenetic variation |
| Epigenetic (histone modifications) | Phylogeny-wide and population level | Assays described in Table S1 | Define a range of histone marks and their regulation across tissues, developmental stages and common-garden physiological manipulations<br>Associate variation in histone marks with all forms of genomic, transcriptomic, proteomic and other classes of epigenetic variation |
| Epigenetic (chromatin biology) | Phylogeny-wide and population level | Assays described in Table S1 | Generate maps of DNA accessibility and define their regulation across tissues, developmental stages and common-garden physiological manipulations<br>Associate changes in chromatin structure with all forms of genomic, transcriptomic, proteomic and other classes of epigenetic variation |
| RNA expression | Phylogeny-wide and population level | RNAseq - potentially stranded protocols (see Table S1) | Define expression of miRNA, mRNA and non-coding RNA across adult tissues, developmental stages and common-garden physiological manipulations <sup>1</sup><br>Associate transcriptomic variation to all forms of genomic, epigenetic and proteomic variation |
| Protein level | Phylogeny-wide and population level | Various possible mass spectrometer platforms – bottom up approach | Define proteome across tissues, developmental stages and common-garden physiological manipulations<br>Associate proteomic variation to all forms of genomic, transcriptomic and epigenetic variation |

## 3. Genome-led science in salmonids: progress, challenges and unresolved questions

Notable progress in understanding of salmonid biology has stemmed from sequencing two salmonid genomes, as well as that of the northern pike *Esox lucius* [12], a sister lineage that did not undergo the salmonid-specific WGD (Fig. 1). Genome-wide analyses have offered key insights into the remodelling and divergence of duplicated genome content and functions during the post-WGD rediploidization process [9, 10]. Population genomics has been revolutionized by genotyping-by-sequencing, whole genome re-sequencing and high-density SNP arrays [13–15], used for example to discover SNPs near the *vgll3* gene that explain 40% of the variation in sea-age at maturity [16, 17], genomic variation explaining the timing of migration [18] and adaptive population differentiation in immune function [19]. Population genomics is now routinely applied in salmonids without a genome sequence, by exploiting conserved synteny with rainbow trout or Atlantic salmon, e.g. [20–23]. Genome-wide approaches have also been applied to improve the accuracy of selection for key production traits (e.g. disease resistance) in breeding programs, either through genomic selection [24–26] or by characterization of major effect loci, e.g. [27, 28]. Further, the salmonid and pike genomes have been used to progress understanding of salmonid phylogeny and species radiation [7] and facilitate characterization of the molecular basis and post-WGD evolution of several physiological systems, including smoltification [29], growth [30], immunity [19, 31, 32] and olfaction [33]. Finally, the recent demonstration of successful genome editing in salmonids [34–37] opens the door for validation of candidate functional genomic elements and causative polymorphisms. Genome editing also has potential to address certain challenges in aquaculture, by creating or introgressing alleles into farmed populations, and by expediting the selection for existing beneficial alleles [38].

Nonetheless, salmonid research and its applications have only just begun to exploit the possibilities of genome-led science. Undoubtedly, a number of unresolved questions and important challenges can be addressed through the FAASG initiative (Box 1).

#### Box 1

The role of functional genome annotation in addressing key challenges for salmonid research and application. Below we list selected key questions, highlight their importance, then briefly describe (*in italics*) how the FAASG initiative will help address them.

##### Aquaculture

What is the functional genetic basis of key performance traits for salmonid aquaculture?

Few causative variants underlying performance trait QTL have been identified. Knowledge of the precise functional variants underpinning QTL will inform the biology of these traits, and facilitate cost-effective selection for favorable alleles.

*Genome annotation is essential to prioritize candidate causal variants. Many traits are influenced by non-coding variants influencing gene expression. The FAASG initiative will aid identification and prioritization of QTL-region variants for key traits.*

How can we optimize genomic selection for genetic improvement in aquaculture breeding programs?

Genomic selection can accelerate genetic gain for traits important to sustainable and profitable aquaculture, such as host resistance to infectious diseases. Predicting breeding values in distant relatives to the training population is challenging, thus necessitating frequent, expensive phenotypic tests.

*The likelihood of SNPs having a functional effect on a trait can be estimated using FAASG functional annotation data. These SNPs can be prioritized in genotyping panels to enable improved prediction accuracy, and persistency of that accuracy, across diverse genetic backgrounds and multiple generations.*

What is the functional genetic basis of recent domestication in salmonid species?

Salmonids are excellent models for the genomic basis of recent domestication, facilitating discovery of genetic variation of importance in adaptation to aquaculture environments. These outcomes can improve hatchery management, health and welfare of farmed fish, and have implications for interactions with wild populations.

*Domestication is likely to have a polygenic basis and be largely due to modification of gene regulation and may also have some epigenetic basis. Functional annotation is essential for researchers to identify sequence and epigenomic variation linked to domestication and response to artificial selection.*

How can genome editing technology contribute to improved aquaculture production?

Genome editing technology, notably CRISPR-Cas9, has potential to enhance aquaculture production directly, by creating or introducing favorable alleles into farmed populations. While regulatory and public acceptance is required, the potential is highlighted by several high profile successes in terrestrial livestock.

*Choosing the correct target to edit is essential, and requires accurate annotation of the reference genome. A function of a SNP, epigenetic mark, non-coding RNA, coding RNA or whole protein can be determined using gene editing. The technology can also be applied to demonstrate causality of variants underlying QTL.*

What is the long term impact of aquaculture escapees on wild populations?

Evaluating and understanding the impacts of aquaculture escapees on wild populations supports risk assessment for the use of native and non-native strains in culture.

*FAASG will improve understanding of the functional differences among populations resulting from genomic variation, and will guide development of tools to effectively track and monitor the genetic impact of escapees on wild populations.*

How can measurement of salmonid health and welfare in aquaculture be improved?

Appropriate biomarkers of stress, health and growth status in salmonid aquaculture are currently difficult to define and far from comprehensive.

*An improved understanding of the genetic and epigenetic regulation of key physiological systems supporting fish health will be guided by the annotated genomes, networks and comparative biology, and will facilitate development of tools to help monitor animal wellbeing in culture.*

##### Ecology, evolution and physiology

What role did the whole genome duplication and subsequent rediploidization play in salmonid evolution and species radiation?

This is a long-standing question of fundamental importance to our understanding of salmonid biology and the role of WGDs in evolution more generally.

*Comparative genomic annotation will improve understanding of how sequence and functional variation arising post-WGD is coupled to trait evolution, including lineage-specific evolution of anadromous life-history and species radiation.*

How important is genetic vs. epigenetic variation in regulating trait variability?

Rapid phenotypic divergence and phenotypic plasticity is a hallmark of many salmonids species, yet remains poorly-characterized. An improved understanding of heritable epigenetic variation and its interaction with both genetic and environmental variation can be exploited in both conservation and aquaculture.

*Functional annotation of epigenetic marks in salmonid genomes, and studies into the role of epigenetic regulation in determining trait variation and phenotypic plasticity are key goals of the FAASG initiative.*

What is the genomic basis of response and adaptation to natural and anthropogenic stressors?

Human-induced environmental changes, including climate change, are already negatively affecting salmonid populations. Understanding the role of genetic and epigenetic variation in physiological response to these changes will be key to predicting, and potentially mitigating, these effects.

*Improved understanding of the functional genomic basis of differential responses to environmental stressors in salmonids may be applied to inform forecasting, mitigation and remedial strategies for challenges associated with climate change*

What role do ‘non-coding’ RNAs have in generating phenotypic variation?

The functions of non-coding RNAs are poorly understood in salmonids. The greater retention of miRNAs in comparison to duplicated genes after WGD suggests important functions in coping with a duplicated genome. Noncoding RNAs may regulate traits of interest to aquaculture and evolutionary biology.

*Comparative functional annotation in salmonids will highlight the location and role of non-coding RNAs in regulation of gene expression and downstream regulation of complex traits.*

How many salmonid species exist, and how can we distinguish them?

The actual number of salmonid species is unknown. Habitat-dependent phenotypes can suggest different species, but genomics and functional genomics methods are ultimately required to answer this question.

*Diverse salmonid species and populations will be targeted in FAASG, providing comparative genome sequences and annotations. This will facilitate development and application of species-specific markers to assess the quantity and diversity of species in the salmonid family.*

## 4. Traits of crosscutting relevance: from aquaculture to evolution (and beyond)

Several traits of importance to aquaculture show extensive natural variation among salmonid species and populations, including disease resistance, growth rate, the control of sexual determination and maturation, and the physiological transition from fresh to saltwater. These traits have crosscutting relevance to multiple scientific fields, both fundamental and applied, and the dissection of their functional genomic architecture under the FAASG initiative will help address challenges faced by the aquaculture sector, along with long-standing research questions. Accordingly, the outcomes of FAASG will facilitate selection of aquaculture strains with improved disease resistance and higher product quality that reach market earlier [39–41], while explaining the evolutionary role of trait variation in wild populations [16, 42–43] and informing management actions influencing population resilience, conservation, and reintroduction [23, 44–46]. Comparing the outcomes of artificial vs. natural selection on functional pathways under different conditions will also help dissect the genetic architecture of traits. For example, different populations will often share genetic variation influencing a trait, but aquaculture and wild conditions impose divergent selective pressures, leading to unique, yet complementary opportunities to understand natural selection and domestication.

## 5. Rationale for linking with FAANG

The FAANG consortium aims to produce comprehensive maps of the functional elements in the genomes of domesticated animal species [11], building on the ENCODE project [47]. Underpinning principles of both consortia include use of robust, standardized experimental protocols based on defined tissues or cell types. These principles apply to both ‘wet lab’ experiments and bioinformatic analyses of data, which provides a comprehensive, reliable, open-access resource for a wide community. The FAASG initiative will link to FAANG, adhere to these principles, and utilise and build on the FAANG protocols and pipelines to avoid redundancy. FAANG is focussed on livestock species with high-quality reference genomes (chicken, pig, cattle and sheep), but with scope for inclusion of other species. The initial focus of FAASG will be the key farmed salmonids (Atlantic salmon and rainbow trout), but will expand to a broader range of lineages of interest to conservation, management and evolution (Fig. 1). In doing so, the initiative will harness wider diversity within a comparative context (section 6) to understand the evolution of functional genome elements following species radiation and WGD. FAASG will provide a FAANG-type model for other lineages with recently-developed genome assemblies, the number of which is rapidly increasing. There will also be great scope for cross-talk between FAASG and research communities for model fish species where functional annotation is advanced, including zebrafish (http://zfin.org/).

## 6. The FAASG framework

The initial approach of FAASG will exploit the full phylogenetic framework of salmonid evolution, documenting functionally important sequence variation and data derived from a core set of experimental assays (section 7) across at nine eight salmonid species (from six out of ten true genera) and northern pike (Fig. 1) and under experimental conditions representative of the traits listed in section 4. This sampling traverses the diversification of salmonid lineages and evolutionary origins of anadromy, a life-history strategy that evolved at least twice independently [48] (Fig. 1) and facilitated species radiations [7]. FAASG will address microevolutionary variation by contrasting wild populations that evolved divergent phenotypes over thousands of years and aquaculture vs wild strains separated by relatively few generations (Fig. 1). This combination of experimental assays and evolutionary analyses done across the salmonid phylogeny (Section 7) will be applied to assess ‘genome function’, thereby addressing a potential shortcoming of the original interpretations of the ENCODE data [49].

## 7. Data and assays

The assays of FAASG are described in Table 1 (also, see Table S1). Annotating distinct classes of sequence variation will identify the genome-wide evolution of orthologous proteincoding genes, along with the large number of retained functional gene duplicates (> 50% of those created) from WGD [9–10]. Population-level sequence variation will inform the role of functional elements in recent phenotypic divergence and adaptation (Table 1). The inclusion of Northern pike (Fig. 1) will enable the ancestral (non-duplicated) state of sequence variation to be inferred, including the direction of divergence between duplicated genes. Comparisons to more distantly related fish with well-annotated genomes, including zebrafish *Danio rerio* [50], three-spined stickleback *Gasterosteus aculeatus* [51], spotted gar *Lepisosteus oculatus* [52], European seabass *Dicentrarchus labrax* [53], and Asian seabass *Lates calcarifer* [54], will allow salmonid-specific changes to be contextualized in the broader framework of teleost evolution, especially with respect to the earlier WGD event in the teleost ancestor.

Transcriptome and proteome phenotypes will be characterized for a panel of tissues and developmental stages, sampled from both sexes under common garden conditions using standardized sampling and analytical protocols that distinguish divergence in expression of duplicated loci [9–10] (e.g. RNA extraction, quality control, library preparation, choice of sequencing platform, and bioinformatic analyses). Discerning the regulation and evolution of transcript complexity (e.g. non-coding, miRNome and splice variants) will necessitate stranded approaches [55] and may be facilitated by capture of full-length transcripts through single molecule real-time sequencing [56]. Standardized proteome expression profiling will also be performed after experimental separation of different cellular fractions.

FAASG will implement genome-wide experimental assays being used or considered under FAANG [11] (Table 1, S1), including: 1) methylation at nucleotide-level resolution (several approaches available, e.g. [57–58]), 2) chromosome accessibility and architecture (via ATACSeq [59]), DNase I footprinting [60], or ChiP-seq approaches), 3) histone modifications (using ChiP-seq approaches [61–62]), 4) genome conformation (via Hi-C [63]) and 5) transcription factor binding occupancy (via ChiP-seq approaches [64]). Many of these assays have yet to be employed in salmonids (Table S1). However, several studies have laid the groundwork for such efforts, and no technical limitations are expected given that these approaches rely on generic techniques and represent conserved features of molecular biology. Initial experiments in Atlantic salmon and rainbow trout will be conducted in the context of regulation across tissues and developmental stages. Assays incorporating different lineages, populations, and physiological manipulations will follow within the wider comparative-phylogenetic framework. Targeted genome editing can subsequently be used to infer causality of sequence variants and functional genomic elements.

## 8. Importance of standardized phenotypic data

Informative genome functional annotation will necessitate standardized measurement and recording of both ecologically and production-relevant traits (section 4) and for the effects of plasticity [65] to be controlled. Comparisons of the genetic architectures for complex phenotypes are confounded not only by the environment in which traits are measured, but also by how those traits are quantified. We view common garden experiments, performed under agreed standardized conditions and treatments, as central to the collection of highquality phenotype data. Salmonids are well-suited for common-garden experiments as they possess external fertilization, high fecundity, and have high survival rates in captivity. In addition, facilities are widely available to raise large numbers of fish under a range of controlled experimental contexts. Such features also facilitate robust and powerful studies to dissect the quantitative genetic basis of complex traits. The standardized recording of both ecologically and production-relevant phenotypes and cataloguing of functional and phenotypic responses, e.g. within the Gene Ontology framework is also high priority. Standardised phenotypic assays will also help interpret the molecular basis of phenotypic variation observed in the numerous wild populations gained by long-term data series, e.g. [66].

## 9. Engaging the research community

The FAASG initiative will promote inclusiveness among all stakeholders and draw in expertise in aquaculture, bioinformatics/biostatistics, genetics, molecular biology, functional genomics, physiology, ecology and conservation, ensuring quality at all levels. A website (http://www.faasg.org/) will report progress, including experimental and computational protocols, publications and datasets, along with contact information for researchers or funders wishing to contribute to efforts. The initiative is being advertised at several scientific conferences (see http://www.faasg.org/) to promote wider awareness.

## Declarations

## List of abbreviations

FAANG: Functional Annotation of Animal Genomes
FAASG: Functional Annotation of All Salmonid Genomes
WGD: Whole genome duplication

## Ethics (and consent to participate)

Not applicable

## Consent to publish

Not applicable

## Competing interests

SB and RJR are respective employees of the Research Council of Norway and Genome British Columbia and made contributions to the scientific writing of this article and the general scientific development of the FAASG initiative.

## Funding

The first FAASG meeting (June 2016, http://www.faasg.org/publications/toronto-workshop-report/) was organized and supported by the International Cooperation to Sequence the Atlantic Salmon Genome (ICSASG). ICSASG funders: The Research Council of Norway (RCN), The Norwegian Seafood Research Fund (FHF), Genome British Columbia (Canada) GBC), The Chilean Economic Development Agency (CORFO) and the Innova Chile Committee (InnovaChile). SB and RJR are respective employees of the Research Council of Norway and Genome British Columbia and made contributions to the scientific writing of this article and the general scientific development of the FAASG initiative.

## Authors’ contributions

Coordinated and drafted the manuscript: DJM, CRP, RDH, BFN. SRS contributed to Figure

1. All authors contributed to manuscript writing and read and approved the final version of this article.

## Availability of data and materials

not applicable

## Acknowledgements

The first FAASG meeting (June 2016, http://www.faasg.org/publications/toronto-workshop-report/) was organized and supported by the International Cooperation to Sequence the Atlantic Salmon Genome (ICSASG). ICSASG funders: The Research Council of Norway (RCN), The Norwegian Seafood Research Fund (FHF), Genome British Columbia (Canada) GBC), The Chilean Economic Development Agency (CORFO) and the Innova Chile Committee (InnovaChile).

---

**Table S1.**
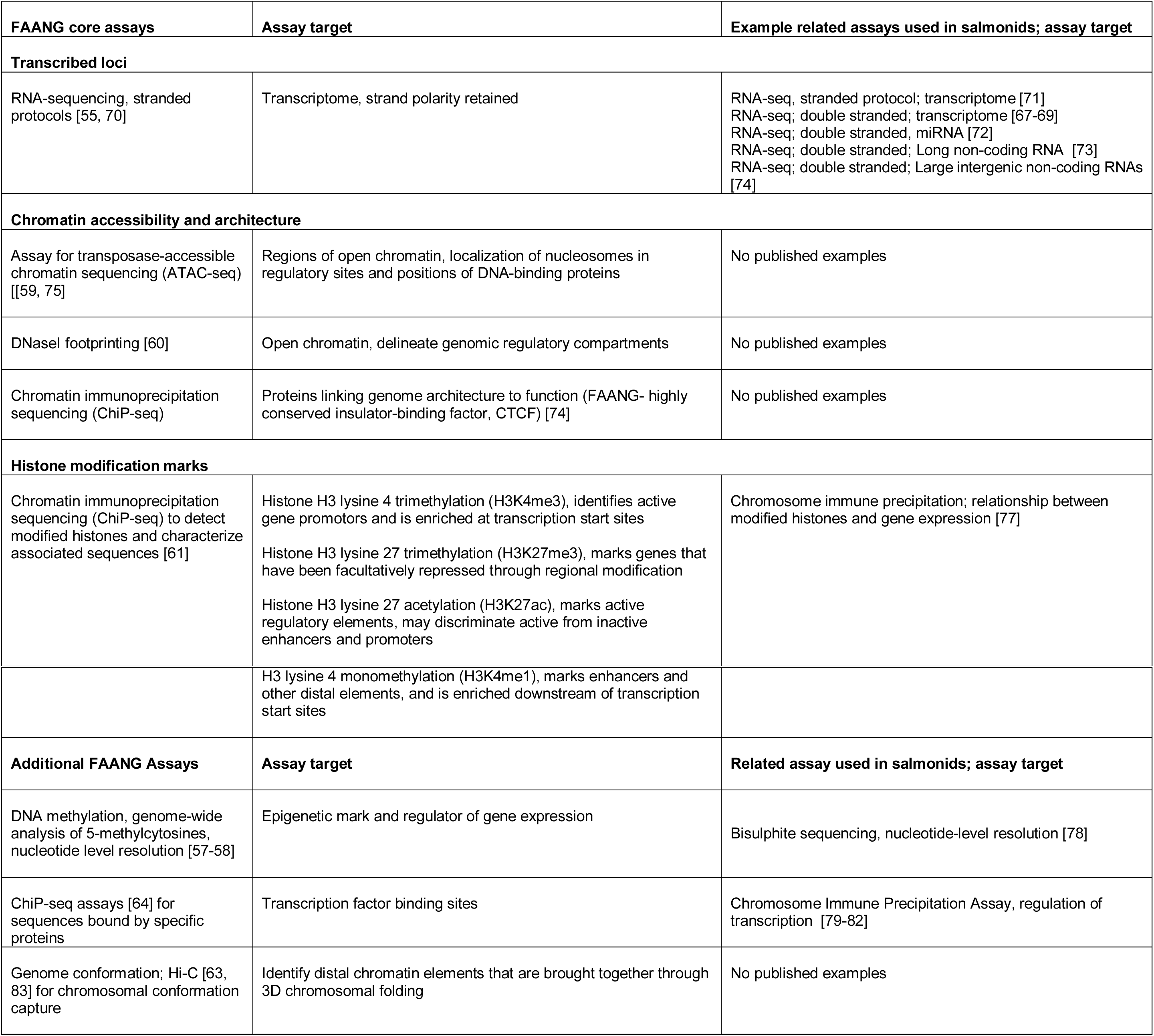
Review of FAANG core/additional assays (as defined in [11]) and relevant work published in salmonids

| **FAANG core assays** | **Assay target** | **Example related assays used in salmonids; assay target** |
| --- | --- | --- |
| **Transcribed loci** |  |  |
| RNA-sequencing, stranded protocols [55, 70] | Transcriptome, strand polarity retained | RNA-seq, stranded protocol; transcriptome [71]<br>RNA-seq; double stranded; transcriptome [67-69]<br>RNA-seq; double stranded, miRNA [72]<br>RNA-seq; double stranded; Long non-coding RNA [73]<br>RNA-seq; double stranded; Large intergenic non-coding RNAs [74] |
| **Chromatin accessibility and architecture** |  |  |
| Assay for transposase-accessible chromatin sequencing (ATAC-seq) [[59, 75] | Regions of open chromatin, localization of nucleosomes in regulatory sites and positions of DNA-binding proteins | No published examples |
| DNaseI footprinting [60] | Open chromatin, delineate genomic regulatory compartments | No published examples |
| Chromatin immunoprecipitation sequencing (ChIP-seq) | Proteins linking genome architecture to function (FAANG- highly conserved insulator-binding factor, CTCF) [74] | No published examples |
| **Histone modification marks** |  |  |
| Chromatin immunoprecipitation sequencing (ChIP-seq) to detect modified histones and characterize associated sequences [61] | Histone H3 lysine 4 trimethylation (H3K4me3), identifies active gene promoters and is enriched at transcription start sites<br>Histone H3 lysine 27 trimethylation (H3K27me3), marks genes that have been facultatively repressed through regional modification<br>Histone H3 lysine 27 acetylation (H3K27ac), marks active regulatory elements, may discriminate active from inactive enhancers and promoters | Chromosome immune precipitation; relationship between modified histones and gene expression [77] |
|  | H3 lysine 4 monomethylation (H3K4me1), marks enhancers and other distal elements, and is enriched downstream of transcription start sites |  |
| **Additional FAANG Assays** | **Assay target** | **Related assay used in salmonids; assay target** |
| DNA methylation, genome-wide analysis of 5-methylcytosines, nucleotide level resolution [57-58] | Epigenetic mark and regulator of gene expression | Bisulphite sequencing, nucleotide-level resolution [78] |
| ChiP-seq assays [64] for sequences bound by specific proteins | Transcription factor binding sites | Chromosome Immune Precipitation Assay, regulation of transcription [79-82] |
| Genome conformation; Hi-C [63, 83] for chromosomal conformation capture | Identify distal chromatin elements that are brought together through 3D chromosomal folding | No published examples |

